# Acid sphingomyelinase inhibition restores RPE homeostasis and photoreceptor function in preclinical Stargardt macular degeneration models

**DOI:** 10.64898/2026.02.20.706984

**Authors:** Colin J. Germer, Sydney Williams, Valencia Fernandes, Ricardo Espinosa Lima, Nilsa R. La Cunza, Li Xuan Tan, Samir Ranjan Panda, Emma Iorio, Fanny M. Elahi, Aparna Lakkaraju

**Author notes:** Correspondence: Aparna Lakkaraju. These authors contributed equally (Colin J. Germer, Sydney Williams).

## Abstract

Stargardt disease, which destroys central high-resolution vision in over 2 million people globally, lacks effective therapies. The primary site of damage in Stargardt disease is the retinal pigment epithelium (RPE), which safeguards photoreceptor health and function. Progressive loss of RPE integrity precedes visual deficits, yet insight into mechanisms driving RPE dysfunction and how this influences disease pathogenesis remains elusive. Here, we addressed this in cell-based and pigmented *Abca4^-/-^* Stargardt mice models using super-resolution imaging, bioinformatics, and biochemical approaches. We show that ceramide accumulation induced by bisretinoid-mediated overactivation of acid sphingomyelinase (ASM) in *Abca4^-/-^* RPE selectively disrupts Rab GTPases and ESCRT machinery involved in apical membrane trafficking and small extracellular vesicle (EV) biogenesis. Consequently, connexin 43 (Cx43) is misrouted from cell-cell junctions into EVs that are released apically by the RPE. This compromises RPE integrity and promotes subretinal immune cell recruitment, leading to photoreceptor dysfunction. Pharmacological ASM inhibition normalizes EV biogenesis and restores Cx43 localization. Decreasing RPE ceramide safeguards RPE structural integrity, limits subretinal microglia, and improves visual function in *Abca4^-/-^* mice. This study underscores the importance of the RPE as a communication hub in the retina and identifies ASM as a potential therapeutic target to prevent progressive vision loss.

## INTRODUCTION

Inherited retinal dystrophies are a leading cause of progressive irreversible vision loss in humans and most are currently untreatable (1). Autosomal recessive Stargardt disease (STGD1), caused by mutations in the ATP-binding cassette A4 (*ABCA4*) gene, is the most common form of inherited macular degeneration, with a prevalence of 1 in 5,000 people. ABCA4 is required for detoxifying vitamin A metabolites formed as by-products of the visual cycle. Loss of ABCA4 function leads to the accelerated formation of toxic bisretinoid adducts in the form of lipofuscin granules in the retinal pigment epithelium (RPE). The RPE, a monolayer of postmitotic polarized epithelial cells that sits between the photoreceptors and the choriocapillaris, performs numerous functions essential for maintaining healthy vision. Pathological accumulation of bisretinoids compromises critical homeostatic functions of the RPE, culminating in RPE atrophy, photoreceptor degeneration, and progressive visual impairment (2–4). There are currently no approved therapies that prevent RPE atrophy or vision loss in Stargardt disease patients (5).

The polarized architecture of the RPE, with distinct apical and basolateral domains demarcated by lateral cell-cell junctions, is critical for its specialized functions. The RPE forms the outer blood-retinal barrier, transports nutrients and metabolites into and out of the retina, participates in phototransduction by recycling vitamin A, and executes daily phagocytosis and removal of photoreceptor outer segment tips (6, 7). Despite the central role of the RPE in the retina, how its structural and functional integrity is maintained over the organismal lifespan remains poorly understood. In other epithelia (e.g., intestine or kidney), a complex and dynamic endosomal network reinforces cellular polarity and communicates with distinct apical and basolateral extracellular environments to coordinate tissue homeostasis (8, 9). Little is currently known about the spatial organization and functions of endosomes in the RPE, or how these are compromised to drive retinal pathology in conditions like Stargardt disease.

Endosome identity is conferred by Rab GTPases, a family of ∼70 ubiquitously expressed proteins that regulate endosome biogenesis, cargo sorting, and transport (10, 11). Defects in specific Rab GTPases and endosome populations are implicated in driving axonal pathology in Alzheimer’s disease (12) and epithelial-mesenchymal transition (EMT) in the colon (13). We have reported that accelerated accumulation of bisretinoids in the RPE of the pigmented *Abca4^-/-^* mice model of Stargardt disease activates acid sphingomyelinase (ASM), the enzyme that hydrolyzes sphingomyelin to ceramide (14). Excess ceramide in the RPE promotes abnormal biogenesis of Rab5+ early endosomes, which facilitate intracellular complement processing and overactivation of the mechanistic target of rapamycin (mTOR), and thus increase RPE susceptibility to metabolic stress (15).

Here, we investigated the contribution of Rab GTPases and polarized membrane trafficking to Stargardt disease pathology. Using *Abca4^-/-^*mice and cell-based models, we report that bisretinoid-induced ceramide accumulation in the RPE specifically impacts Rab GTPases Rab11a, Rab27a, and Rab37, which regulate apical membrane trafficking, exocytosis, and junctional integrity. Further, ceramide drives inward budding of endosomal limiting membranes, leading to increased biogenesis of intraluminal vesicles (ILVs). These abnormalities in membrane trafficking induce mis-sorting of the gap junction protein connexin 43 (Cx43) into ILVs that are released apically as small extracellular vesicles (EVs) by the RPE, towards the neural retina. This compromises RPE structural integrity and induces the migration of immune cells to the subretinal space, culminating in photoreceptor dysfunction. The tricyclic antidepressant desipramine, a functional ASM inhibitor that decreases RPE ceramide (14, 16), normalizes Rab GTPase expression, restores Cx43 localization, limits microglial migration, and improves photoreceptor function in *Abca4^-/-^* mice.

Structural remodeling of the RPE is a feature of many blinding diseases including Stargardt disease and age-related macular degeneration (AMD), but the underlying mechanisms and its connection to other pathological features have been unclear. This study demonstrates that disruption of specific endosomal populations due to overactive ASM not only compromises RPE homeostasis but also drives immune cell activation and photoreceptor dysfunction in the retina.

## Results

### Apical recycling and EV biogenesis machinery are specifically dysregulated in Abca4^-/-^ mice RPE

To investigate membrane trafficking pathways in healthy and diseased RPE, we first evaluated the expression and subcellular localization of Rab GTPases by immunofluorescence imaging and immunoblotting. Our data revealed increased expression and predominantly apical localization of Rab11a and Rab37 in 10-month-old *Abca4^-/-^* mice RPE compared to age-matched wildtypes (**Figures 1A and 1B**). We next asked whether this was indicative of a global disruption of Rab GTPases or selective for specific membrane trafficking routes. Immunoblotting data showed that *Abca4^-/-^* RPE had significantly increased expression of Rab11a, which directs apical recycling endosomes, and Rab27a and Rab37, which regulate apical exocytosis, whereas other recycling Rabs (Rab8 and Rab10) and the late endosomal Rab7 were unchanged (**Figures 1C and 1D**).

**Figure 1.**
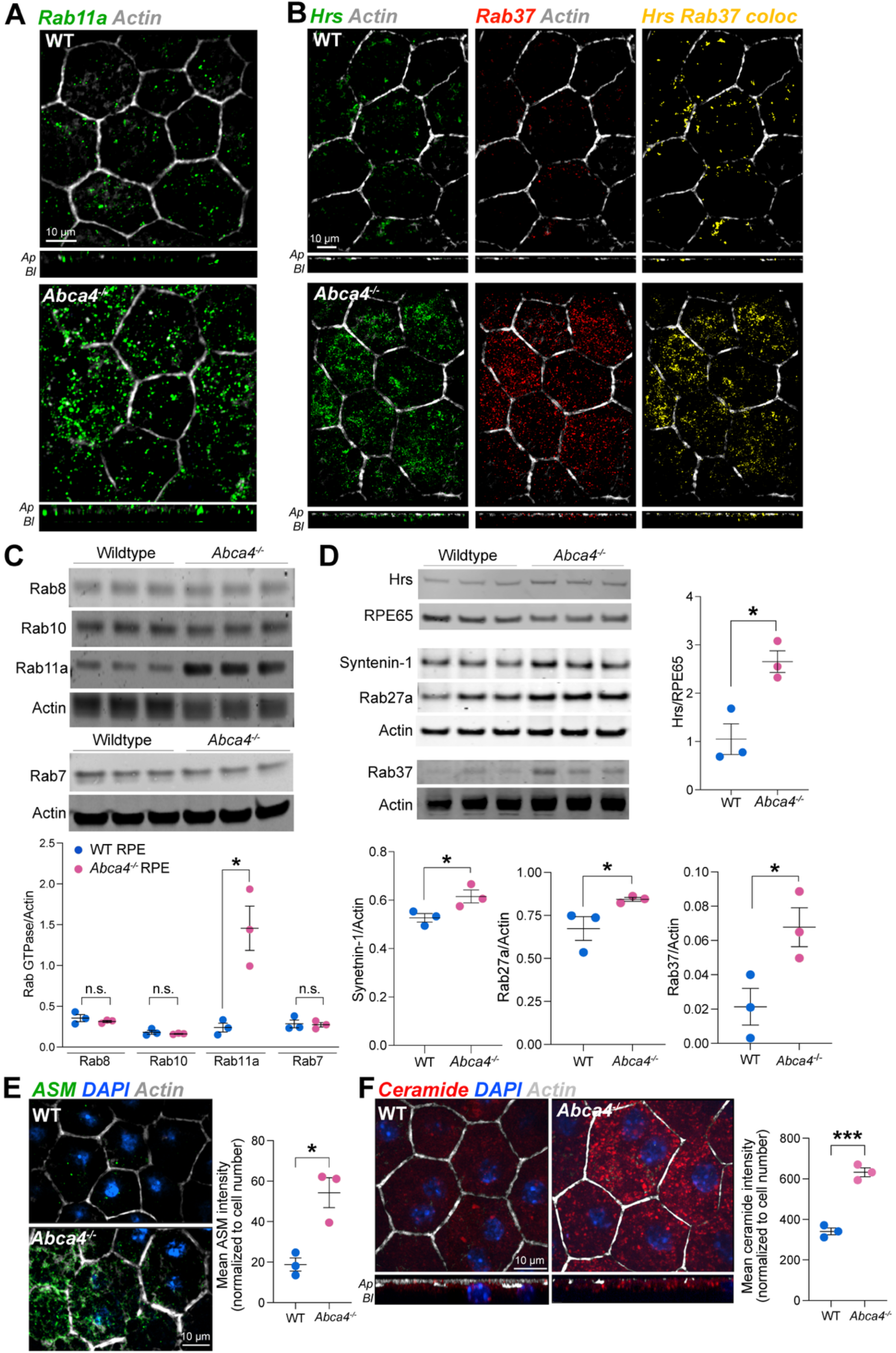
Disruption of apical trafficking and exosome biogenesis machinery in *Abca4^-/-^* RPE. Immunostaining for **(A)** Rab11 and **(B)** Rab37 (red) and Hrs (green) in RPE flatmounts from 10-month-old wildtype and *Abca4^-/-^* mice. Colocalization between Hrs and Rab37 is shown in yellow. Cell boundaries are demarcated with the actin stain phalloidin (grey). **(C)** Immunoblots and quantification of Rab8, Rab10, Rab11, and Rab7 in RPE lysates from wildtype and *Abca4^-/-^* mice. Mean ± SEM *, p < 0.05, n = 3 mice per genotype; each lane denotes one biological replicate. n.s. - not significant. **(D)** Immunoblots and quantification of Hrs, Syntenin-1, Rab27a, and Rab37 in mice RPE lysates. Mean ± SEM *, p < 0.05, n = 3 mice per genotype. **(E)** ASM (green) and (F) Ceramide (red) immunostaining and quantification in mice RPE flatmounts. Mean ± SEM *, p < 0.05, ***, p < 0.0001, n = 3 mice per genotype.

As Rab11a, Rab27a, and Rab37 also modulate the secretion of EVs (17–19), we investigated other EV biogenesis machinery in *Abca4^-/-^* RPE. The endosomal sorting complex required for transport-0 (ESCRT-0) protein Hrs, and the syndecan-binding protein syntenin-1, which regulate cargo sorting and biogenesis of intraluminal vesicles (ILVs), were upregulated in *Abca4^-/-^* RPE (**Figure 1D**). EV biogenesis is regulated by multiple players, including ceramide, a cone-shaped lipid that induces negative membrane curvature to generate ILVs (20, 21). Compared to age-matched wildtype mice RPE, *Abca4^-/-^* RPE had significantly more ASM and, consequently, more ceramide, which was concentrated at the apical surface (**Figures 1E and 1F**). The increase in ASM levels was likely due to cholesterol-mediated sequestration of the ASM cofactor, bis(monoacylglycerol)phosphate (BMP), which stabilizes ASM and prevents its lysosomal degradation in *Abca4^-/-^* RPE as we have previously reported (14).

Taken together, these data indicate specific disruption of machinery that regulates apical membrane trafficking and EV biogenesis/secretion in *Abca4^-/-^* RPE. This, coupled with the apical accumulation of ceramide, could promote abnormal release of EVs towards the photoreceptors by diseased RPE.

### ASM overactivation drives biogenesis and apical secretion of EVs from polarized RPE

Several distinct mechanisms and machinery function in EV biogenesis and secretion: ESCRT proteins, which participate in membrane remodeling and cargo recruitment; Syntenin-1, which is thought to regulate an ESCRT-independent pathway; ceramide-mediated inward budding of ILVs; and lastly, Rab11a, Rab27a, and Rab37, which have been shown to specifically induce apical EV secretion in polarized epithelia (17, 18, 22). Based on our observations that all three major pathways/machineries are impacted in *Abca4^-/-^* RPE, we first evaluated EV biogenesis and secretion using a well-characterized *in vitro* model of primary polarized RPE (23–25). To simulate Stargardt disease, primary porcine RPE cultured on semi-permeable membrane supports were treated with the lipofuscin bisretinoid A2E to achieve intracellular concentrations comparable to those in *Abca4^-/-^* RPE and Stargardt disease patients (14). EVs released into the apical or basolateral media were purified according to published protocols (26, 27) (**Figure 2A**).

**Figure 2.**
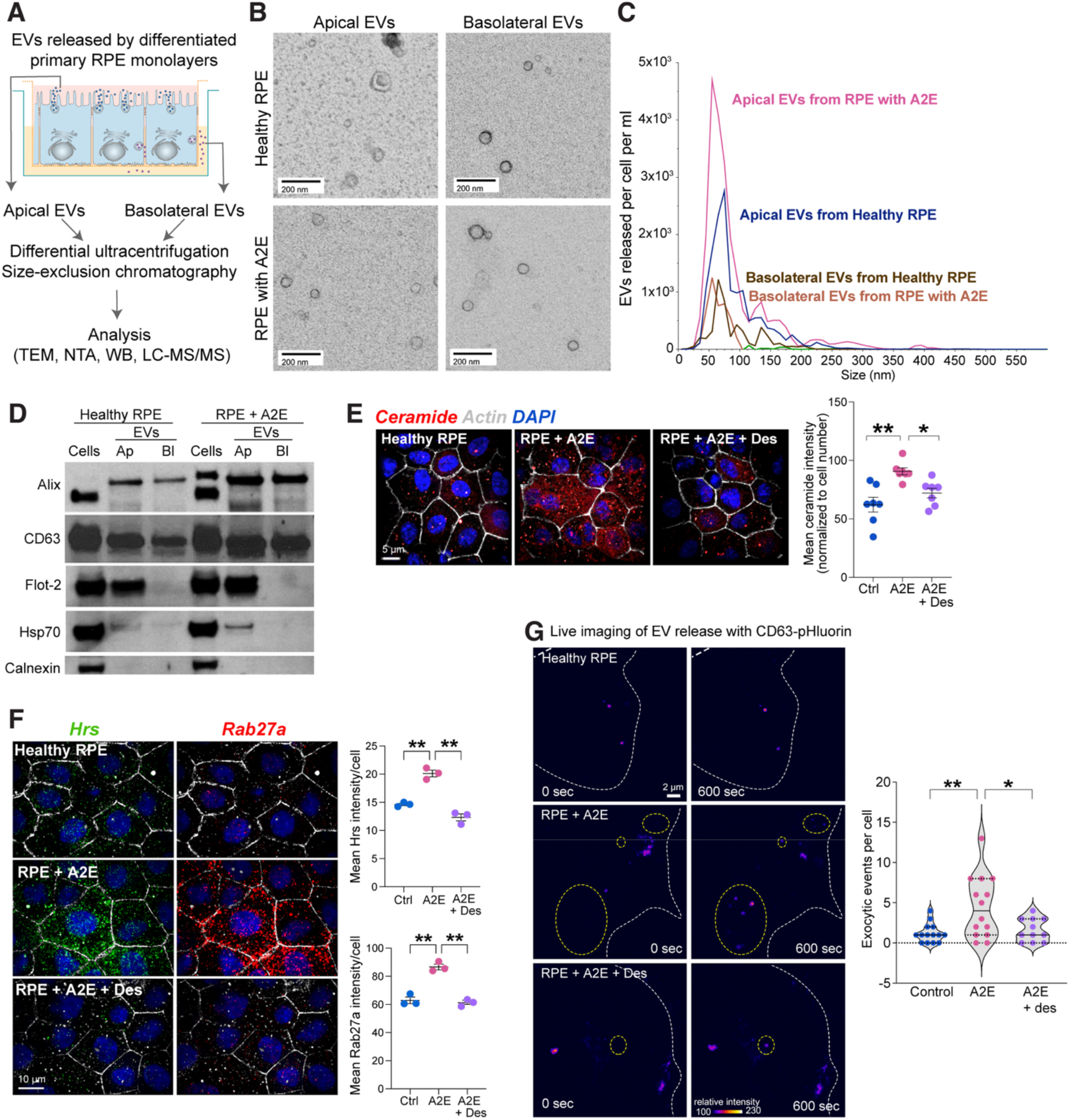
Ceramide increases the biogenesis and apical secretion of extracellular vesicles from differentiated polarized primary RPE monolayers. **(A)** Protocol used to purify EVs from apical and basal media of differentiated primary RPE monolayers cultured on semipermeable Transwell filters. **(B)** Representative transmission electron micrographs of EVs secreted into the apical and basolateral media by healthy RPE cultures or RPE with A2E. **(C)** Number and size distribution of apical and basolateral EVs secreted by healthy RPE cultures or RPE with A2E measured by nanotracking analysis. **(D)** Representative immunoblot of total cell lysates (Cells), and EVs released apically (Ap) or basolaterally (Bl) from polarized RPE cultures with or without A2E. **(E)** Ceramide immunostaining (red) and quantification in healthy RPE, RPE with A2E alone or after treatment with desipramine (Des, 10 µM, 3 h). **(F)** Hrs (green) and Rab27a (red) immunostaining in healthy RPE, RPE with A2E alone, or after desipramine treatment. In E and F, nuclei are stained with DAPI (blue) and cell boundaries are demarcated by the actin stain phalloidin (white). Mean ± SEM *, p <0.05, **, p <0.01; n = 3 independent experiments, with ∼300 cells analyzed per experiment. **(G)** Stills from live imaging of CD63-pHluorin exocytosis (yellow circles) in primary RPE cultures with A2E alone or after treatment with desipramine. Cell boundaries: dashed white lines. Violin plot shows number of exocytic events observed per cell in 600 seconds. Mean ± SEM, *, p < 0.001, n = 12-14 cells per condition.

Negative stain transmission electron microscopy (TEM) showed characteristic vesicles with a size range of 50-100 nm for both apical and basolateral EVs released by healthy RPE and RPE with A2E (**Figure 2B**). We next measured the number and size distribution of these EVs using nanoparticle tracking analysis (NTA). These data confirmed polarized secretion of EVs by the RPE, with more EVs released into the apical media, in agreement with published reports that measured EV secretion from polarized primary porcine RPE monolayers and iPSC-RPE cultures (24, 28–30). RPE with A2E secreted ∼2-fold more EVs/cell/ml into the apical media compared to healthy RPE, whereas the number of EVs released into the basolateral media was unchanged (**Figure 2C**). Apical EVs released by RPE with A2E had a smaller and narrower size distribution (peak diameter 55 nm vs 75 nm for healthy RPE) compared to those secreted by healthy RPE, likely because increased Hrs could drive the formation of uniformly sized small ILVs in RPE with A2E as has been reported in HeLa cells (31). Immunoblotting confirmed that RPE-derived EVs were enriched in canonical EV proteins such as CD63, flotillin-2, and Hsp70, whereas other organelle markers such as the ER resident protein calnexin were absent, indicating the purity of our EV preparations (**Figure 2D**).

As ESCRT machinery, syntenin-1, and ceramide have been implicated in distinct EV biogenesis pathways, we next sought to establish which of these was predominantly responsible for aberrant apical EV secretion in diseased RPE. We focused on ceramide because, apart from inducing negative membrane curvature, it can also modulate membrane localization and function of recycling Rab GTPases such as Rab11 (32, 33). We first sought to determine if ceramide promotes apical EV secretion from other polarized epithelia as a general mechanism. We measured EVs released into the apical and basal media by polarized Madin-Darby canine kidney (MDCK) monolayers. To induce ceramide accumulation independent of A2E, we used U18666A (U18), which increases ceramide via cholesterol-mediated activation of ASM like A2E (**Figure S1A**) (16). TEM, NTA, and immunoblotting analyses confirmed that U18 significantly increased apical secretion of EVs by MDCK cells (**Figure S1B-D**).

Next, to modulate ceramide levels in the RPE, we used the tricyclic antidepressant desipramine, which we previously reported inhibits ASM and decreases ceramide in RPE with A2E and in *Abca4^-/-^*RPE (14–16). Desipramine also decreases ceramide-induced mTOR activation, which could indirectly modulate Hrs as mTOR has been reported to stabilize Hrs against proteasomal degradation (15, 34). A short (3 h) treatment with desipramine decreased ceramide and restored Hrs and Rab27a levels and expression comparable to those in healthy RPE (**Figures 2E and 2F**). These data indicate that ceramide acts upstream of Rabs and ESCRT machinery in regulating EV biogenesis and secretion by the RPE.

To follow EV dynamics in RPE expressing CD63 tagged with a pH-sensitive GFP (CD63-pHluorin), we used high-speed live cell imaging. The GFP fluorescence of this reporter is pH-sensitive and quenched in the acidic pH of MVBs. Upon fusion of MVBs with the plasma membrane, the neutral pH of the extracellular milieu allows GFP to fluoresce and is indicative of EV release (35). In time-lapse imaging over 10 minutes, we observed ∼2-3-fold more EV release events in RPE with A2E compared to healthy RPE whereas desipramine significantly decreased exocytic events in RPE with A2E (**Figure 2G**). These data support a model where excess RPE ceramide due to ASM overactivation remodels specific trafficking pathways that drive apical EV secretion in diseased RPE.

### The gap junction protein connexin 43 (Cx43) is missorted into apical EVs in diseased RPE

To establish the physiological relevance of RPE-derived EVs, we used LC-MS/MS to determine the protein cargo of EVs secreted by healthy RPE cultures or RPE with excess ceramide. Proteomic analysis identified 170 proteins, with 107 common proteins, 5 unique to healthy RPE, and 58 unique proteins sorted into EVs in stressed RPE (**Figure S2A**). These included the tetraspanins CD9 and CD81, and proteins recently identified as core EV components such as syntenin-1 and ADAM10 (36–38) (**Table S1**). Analysis of protein-protein interactions (PPI) and gene ontology (GO) mapping showed significant enrichment of processes that regulate exosome biogenesis, retina homeostasis, and cell-cell junction assembly (**Figures S2B and S2C**). Of the differentially sorted cargo, the gap junction protein connexin 43 (Cx43) was highly enriched (∼1,000-fold) in EVs from RPE with increased ceramide compared to EVs released by healthy RPE.

To confirm the presence of Cx43 in EVs secreted by diseased RPE, we performed super-resolution imaging of single EVs. Purified apical EVs released by healthy RPE or RPE with A2E were triple labeled with antibodies to CD9, ceramide, and Cx43. Direct stochastic optical reconstruction microscopy (dSTORM), which allows imaging of protein cargo in individual EVs (39, 40), and quantification of colocalization revealed that a significant number of CD9+ EVs released by RPE with A2E also contained Cx43 and ceramide, whereas EVs from healthy RPE were largely devoid of Cx43 (**Figure 3A**).

**Figure 3.**
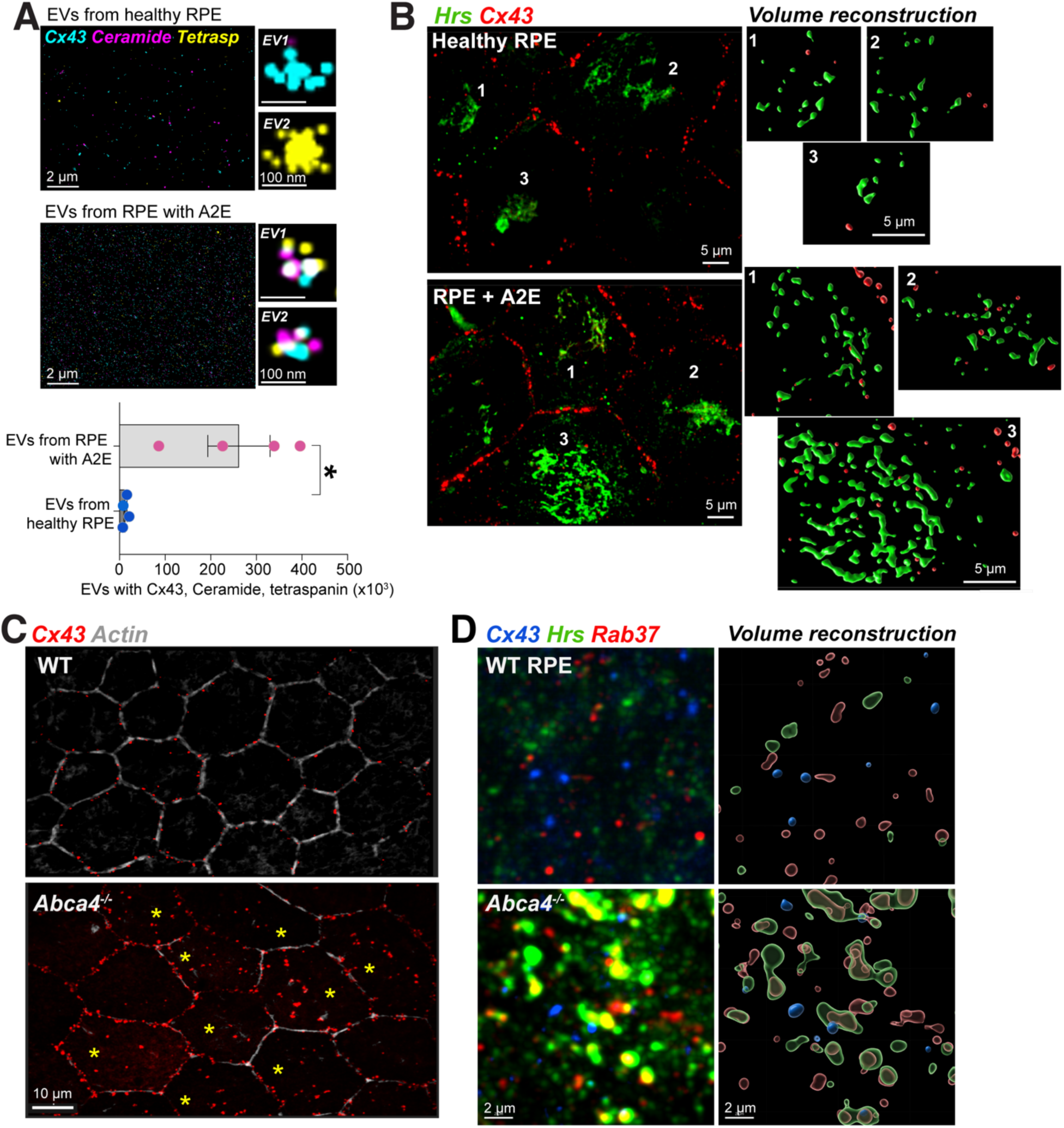
Connexin 43 is sorted into EVs released by diseased RPE. **(A)** Representative dSTORM super-resolution microscopy images showing presence or absence of Cx43 (blue) and ceramide (pink) in tetraspanin (yellow) containing EVs released by healthy RPE or RPE with A2E at single-particle level. Right panels: High-magnification images of two representative EVs from healthy and A2E-treated RPE. Number of EVs containing Cx43, ceramide, and tetraspanin was quantified using CODI software (ONI). Mean ± SEM, n = 4 samples per condition. *, p <0.05. **(B)** Primary RPE cultures (healthy or with A2E) stained for Hrs (green) and Cx43 (red). Right panels: high-magnification volume reconstructions of Hrs and Cx43 in the indicated cells on the left. **(C)** Cx43 (red) immunostaining in 18-mo wildtype and *Abca4^-/-^* mice RPE flatmounts. Yellow asterisks indicate cells with intracellular Cx43. **(D)** High-magnification images of Cx43 (blue), Hrs (green) and Rab37 (red) immunostaining in 18-mo wildtype and *Abca4^-/-^* mice RPE flatmounts. Right panels: volume reconstructions showing association of Cx43, Hrs, and Rab37.

Residence of Cx43 on the membrane is tightly regulated by phosphorylation and ubiquitylation, which direct Cx43 endocytosis and intracellular trafficking, respectively (41, 42). Once internalized, Cx43 associates with ubiquitin-binding ESCRT proteins such as Hrs and Tsg101, enabling transport to multivesicular late endosomes for either lysosomal degradation (42, 43) or sorting into EVs (44). Immunostaining and volume reconstruction showed intracellular Cx43 puncta in association with Hrs in RPE with A2E, but not in healthy RPE monolayers (**Figure 3B**). This pattern was replicated *in vivo*: in contrast to the defined membrane localization of Cx43 in wildtype mice RPE, we observed significant levels of intracellular Cx43 in 18-month-old *Abca4^-/-^* mice (**Figure 3C**; also see **Figure 5E**). Super-resolution imaging showed increased association of intracellular Cx43 with Hrs and Rab37 in *Abca4^-/-^*RPE (**Figure 3D**).

Given that Rab37 has been implicated in apical secretion of EVs from polarized epithelia (18), these data suggest that increased Hrs, Rab37, and ceramide in *Abca4^-/-^* RPE coordinate sorting of Cx43 into late endosomal ILVs that are destined for secretion as EVs rather than lysosomal degradation.

### Sub-retinal immune cells in Abca4^-/-^ mice are enriched in Cx43

Cx43-containing EVs have been reported to induce cell migration, ECM remodeling, senescence, and inflammation in diverse cell types such as chondrocytes, endothelial cells, and cardiomyocytes (45–47). The specific enrichment of Cx43 in EVs released apically by RPE with A2E led us to ask whether these EVs could direct migration of retinal microglia from their homeostatic niches in the inner and outer plexiform layers into the subretinal space. Immunostaining of 18-month-old RPE flatmounts revealed the presence of cells expressing IBA1 (ionized calcium binding adaptor molecule), CD68, and TMEM119, markers for activated retinal microglia (48, 49), in the subretinal space between photoreceptors and the RPE in *Abca4^-/-^* mice (**Figures 4A**, **4B, and S3**). The location of subretinal microglia corresponded with empty or “bald” spots of actin staining in the RPE monolayer, and surrounding RPE appeared dysmorphic or enlarged, indicative of apical membrane remodeling and compromised RPE structural integrity in aged *Abca4^-/-^* mice (**Figures 4A and 4C**, yellow squares). The presence of numerous rhodopsin-positive phagosomes (**Figure 4C**) suggested that these subretinal microglia were phagocytic and potentially pro-inflammatory (50, 51).

**Figure 4.**
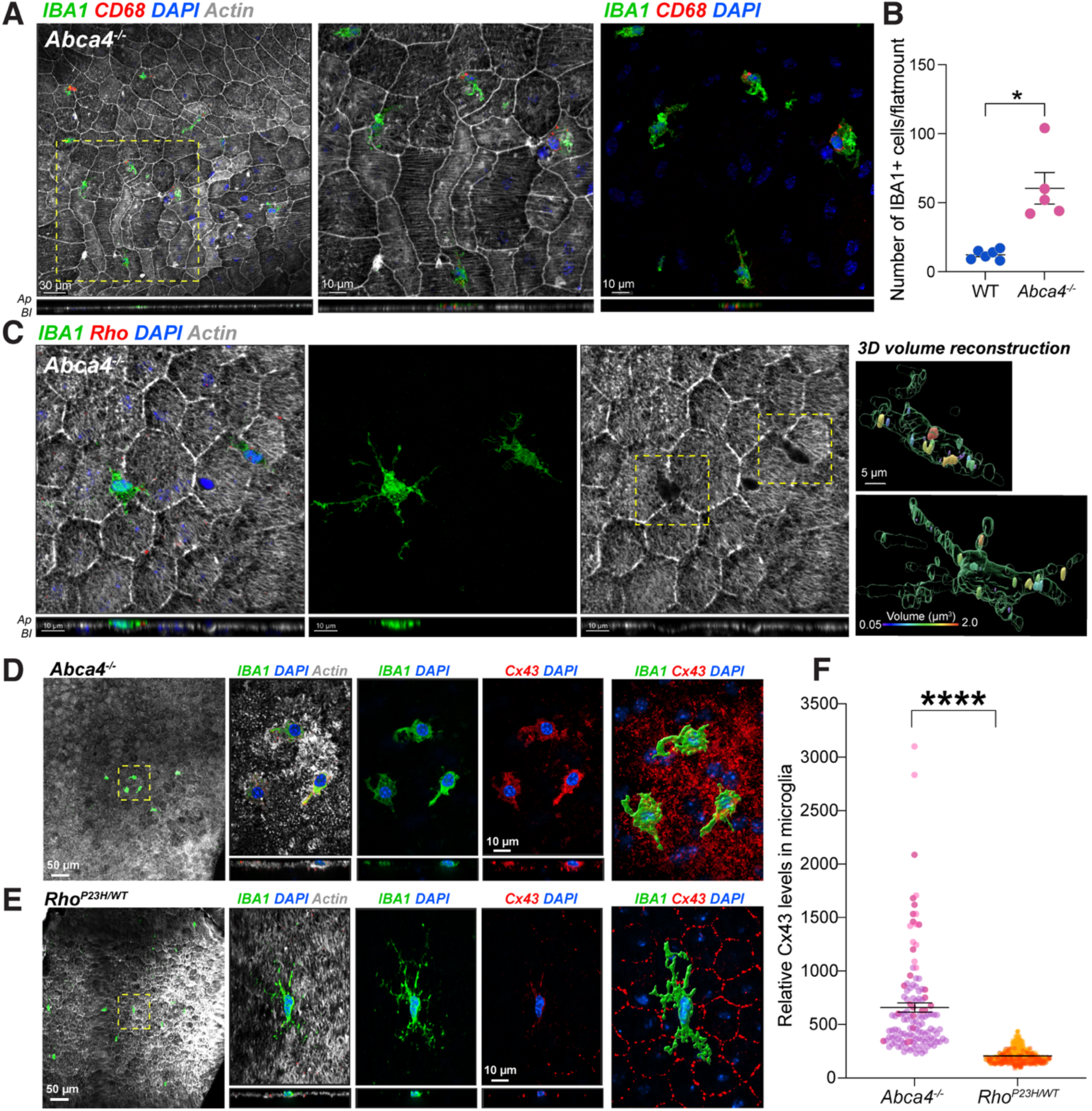
Subretinal immune cells in *Abca4^-/-^* mice are enriched in connexin 43. **(A)** Immunostaining of 18-mo *Abca4^-/-^* mice RPE flatmounts shows IBA1 (green) and CD68 (red) expressing immune cells in the subretinal space. **(B)** Number of subretinal IBA1+ cells per flatmount in 18-month-old WT or *Abca4^-/-^*mice. Mean ± SEM, n = 5-6 mice per group. *, p <0.05. **(C)** IBA1+ (green) microglia contain rhodopsin (red) and correspond to “bald patches” (yellow squares) on the RPE revealed by lack of actin staining (gray). 3D volume reconstructions showing phagosomes within microglia in *Abca4^-/-^* mice. Color scale indicates volume of phagosomes. **(D)** IBA1-positive subretinal microglia in Abca4-/- mice are enriched in Cx43 (red). Rightmost panel shows volume reconstruction of microglia sitting atop RPE with disrupted Cx43 organization. **(E)** IBA1-positive subretinal microglia in Rho*^P23H/+^* mice have barely detectable Cx43 (red). Rightmost panel shows volume reconstruction of microglia sitting atop the RPE with intact Cx43 organization. **(F)** Superplot shows relative levels of Cx43 in subretinal microglia of *Abca4^-/-^* and Rho*^P23H/+^* mice. A total of123 subretinal IBA1+ cells from *Abca4^-/-^* mice and 943 subretinal IBA1+ cells from Rho*^P23H/+^* mice were analyzed. Mean ± SEM, n = 3 mice per group. ****, p <0.0001.

Next, we asked whether Cx43-EVs released by diseased RPE could act as a chemoattractant to induce microglial migration. We used an indirect approach to circumvent inherent challenges associated with accurately tracking the fate and function of RPE-derived EVs *in vivo* in the retina (52). We reasoned that if RPE-derived Cx43-EVs are responsible for microglial recruitment, these subretinal microglia should have higher levels of Cx43 compared to microglia in diseases where retinal pathology is not driven by the RPE. To test this hypothesis, we compared Cx43 levels in subretinal microglia in *Abca4^-/-^* mice with those in Rho*^P23H+^* mice, a model for autosomal dominant retinitis pigmentosa. The P23H mutation causes rhodopsin aggregation within photoreceptors, leading to progressive photoreceptor degeneration accompanied by microglial infiltration into the subretinal space, although the RPE remains relatively unaffected (53). Our data showed that IBA1+ subretinal cells in *Abca4^-/-^* subretinal microglia were enriched in Cx43, in contrast to the barely detectable Cx43 in Rho*^P23H/+^* subretinal microglia (**Figures 4D**–**4F**). Furthermore, Cx43 localization in Rho*^P23H/+^* RPE was restricted to cell-cell junctions comparable to its organization in wildtype mice, in contrast to the disorganized and increased intracellular localization of Cx43 in *Abca4^-/-^* RPE. Altogether, these data suggest that Cx43-EVs released by diseased RPE could induce RPE structural remodeling and microglial recruitment.

### The functional ASM inhibitor desipramine decreases ceramide and EV biogenesis in Abca4^-/-^ mice RPE

To directly evaluate the role of RPE-derived EVs in driving retinal pathology in *Abca4^-/-^* mice, we treated mice with desipramine to modulate EV biogenesis and secretion *in vivo*. Compared to vehicle-treated mice, desipramine (10 mg/kg administered intraperitoneally, 3 times/week for 8 weeks) decreased RPE ceramide in 18-month-old *Abca4^-/-^* mice comparable to wildtype levels (**Figure 5A**). Quantification of MVBs and ILVs using TEM showed no difference in the number, size, or subcellular distribution of MVBs between wildtype and *Abca4^-/-^* RPE; however, the number of ILVs per MVB, which was significantly higher in *Abca4^-/-^* RPE decreased in response to desipramine treatment (**Figures 5B**, **5C, S4A, and S4B**). These data not only confirm our *in vitro* studies but also underscore a central role for ceramide in the biogenesis of EVs in the RPE.

**Figure 5.**
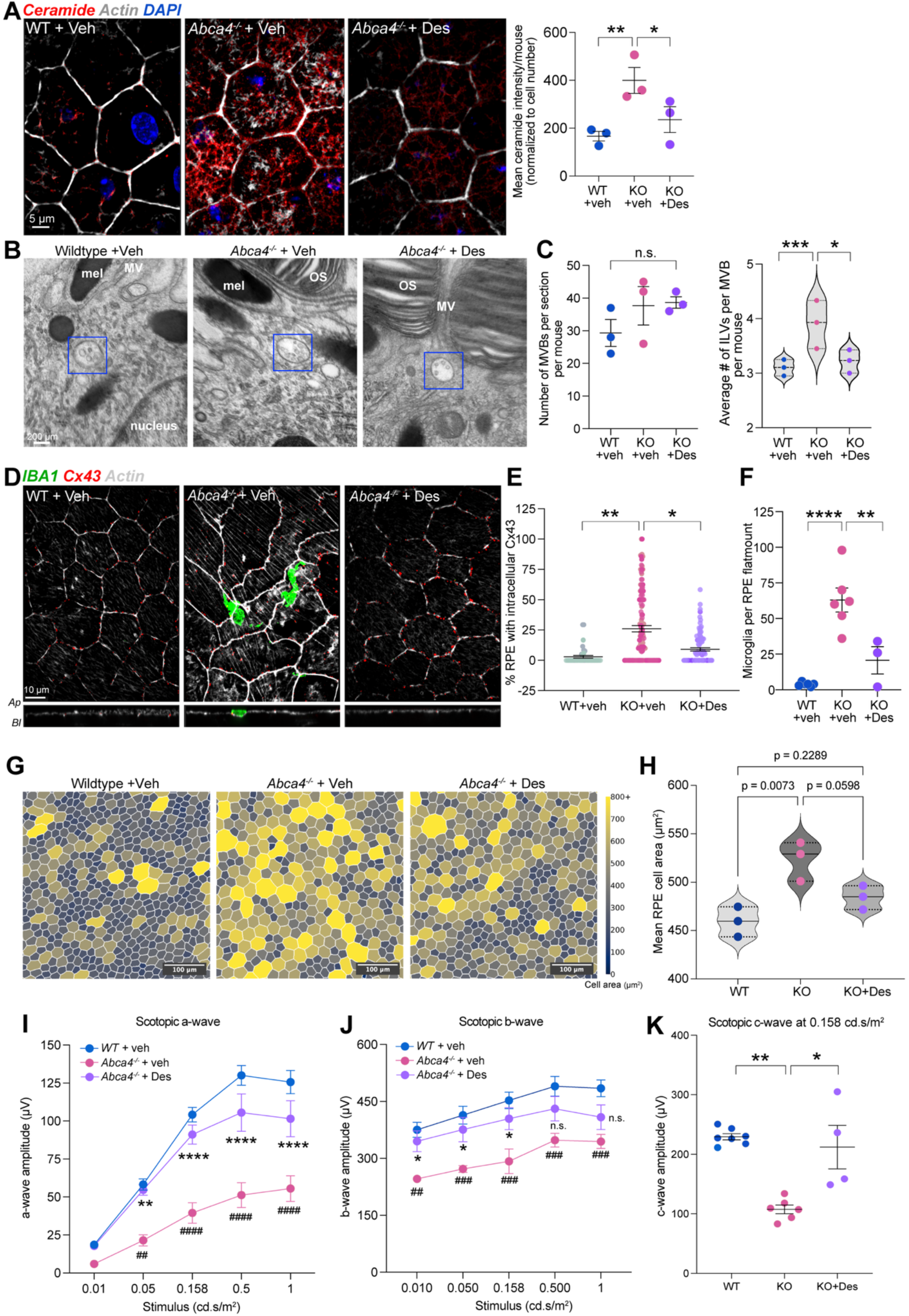
Decreasing RPE ceramide normalizes EV biogenesis, prevents microglial infiltration, and improves RPE and photoreceptor function in aged *Abca4^-/-^* mice. **(A)** Ceramide immunostaining (red) and quantification of RPE flatmounts from 18-mo mice treated with vehicle or desipramine (10 mg/kg, i.p., 3 times/week for 8 weeks). Mean ± SEM, n = 3 mice per group. *, p <0.05; **, p < 0.005. **(B)** Representative TEMs of RPE from vehicle treated wildtype and *Abca4^-/-^* mice or desipramine-treated *Abca4^-/-^* mice. Multivesicular bodies (MVBs) containing intraluminal vesicles (ILVs) are denoted by blue squares. OS: outer segments, Mel: melanosomes, MV: microvilli. Scale bar: 200 nm. **(C)** Number of MVBs and ILVs per MVB in wild-type and Abca4-/- mice RPE. ∼45 MVBs per mouse were analyzed, n = 3 mice per group. **, p < 0.005; ***, p < 0.001. **(D)** Representative images of RPE flatmounts stained for IBA1 (green) and Cx43 (red) in mice treated with vehicle or desipramine. **(E)** Percent of RPE cells with intracellular Cx43 and **(F)** IBA1-positive sub-retinal cells in RPE flatmounts from mice treated with vehicle or desipramine. Mean ± SEM, n = 3-6 mice per group. **, p < 0.005, ****, p <0.0001; one-way ANOVA, Holm-Sidak’s multiple comparison tests. **(G)** Representative images from morphometric analyses of the RPE cell sheet in 18-month-old mice. **(H)** Violin plots of mean RPE cell area (µm^2^) in18-month old mice. 3 mice per genotype/treatment. The entire flatmount was analyzed for each mouse. **(I-K)** Scotopic a-, b-, and c-waves in 18-mo wildtype and *Abca4^-/-^* mice treated with vehicle or desipramine. Mean ± SEM, n = 4-7 mice per group. Vehicle-treated *Abca4^-/-^*mice significantly different from corresponding WTs: ##, p < 0.01; ###, p < 0.001; ####, p < 0.0001. Vehicle-treated *Abca4^-/-^* mice significantly different from desipramine-treated *Abca4^-/-^* mice, *, p<0.05; **, p < 0.01; ****, p < 0.0001.

### ASM inhibition restores RPE integrity, limits subretinal microglia, and improves visual function in Abca4^-/-^ mice

After establishing that desipramine normalizes EV biogenesis in the RPE *in vivo*, we next investigated its impact on key aspects of EV-related retinal pathology in 18-month-old *Abca4^-/-^* mice. Our data show that desipramine restored Cx43 localization to the RPE cell membrane and limited microglial migration to the subretinal space in *Abca4^-/-^* mice (**Figures 5D**–**5F**). Morphometric analyses of the RPE cell sheet revealed a significantly larger mean cell area in *Abca4^-/-^* mice, which decreased after desipramine treatment (**Figures 5G and 5H**).

Pigmented *Abca4^-/-^* mice develop progressive visual deficits after 18 months when assessed by scotopic electroretinograms (ERGs) (54) (**Figure S4C**). Desipramine treatment significantly enhanced a-wave and c-wave amplitudes indicating improved photoreceptor and RPE function and had a moderate impact on b-wave amplitudes, which reflect the function of the inner retina (**Figures 5I**–**5K**). These studies suggest that therapeutic approaches that normalize ASM activity and ceramide levels in the RPE could be beneficial in restoring RPE homeostasis and photoreceptor function in Stargardt disease.

## DISCUSSION

Progressive loss of RPE integrity and function are characteristic features of Stargardt disease (55), yet insight into the underlying molecular mechanisms and how it relates to immune cell activation and photoreceptor dysfunction remains unclear. Here, we show that specific disruption of apical membrane trafficking in the RPE initiates a cascade that encompasses multiple aspects of disease pathology. Selective disruption of Rab GTPases and ESCRT proteins in diseased RPE promotes the biogenesis and apical secretion of EVs containing Cx43. Intracellular mis-sorting of Cx43 towards the EV route in aged *Abca4^-/-^* mice temporally tracks with RPE structural changes, the appearance of activated microglia in the subretinal space, and progression of photoreceptor functional deficits. By demonstrating that ASM inhibition even relatively late in this cascade halts or reverses these sequelae, our data establish RPE ceramide as a central driver of Stargardt disease pathology.

Rab GTPases are critical regulators of cell fate as they control organelle biogenesis, trafficking, and function (10, 56). Alterations in Rab GTPase function due to mutations in the Rab Escort Protein 1 (REP1) cause choroideremia, an inherited retinal degeneration characterized by deficits in organelle function in the RPE and eventual central vision loss (57, 58), suggesting that the RPE is uniquely sensitive to disruptions in Rab GTPase activities. A key finding in this study is the selective upregulation of Rabs 11a, 27a, and 37 in Stargardt disease models. However, apart from a role for Rab27a in melanosome transport (59), little is known about the functions of these Rabs in the RPE. Studies in other tissues suggest that they regulate junctional stability, cell polarity, and cancer: Rab11 interferes with E-cadherin recycling from cell-cell junctions in keratinocytes and binds Cx43 to direct ciliogenesis in the frog epidermis, whereas Rab37-mediated exocytosis promotes tumor metastasis (13, 60–62). Our studies show that in the RPE, these Rabs redirect Cx43 away from lysosomal degradation and promote its sorting into EVs destined for apical release.

*In vitro* studies using primary porcine RPE cultures and iPSC-RPE have reported both homeostatic and stress-induced EV secretion with specific cargo sorted into EVs destined for release at the apical or basal domains (24, 28, 29, 63, 64). While these studies support our observation of preferential apical EV secretion by the RPE, the molecular machinery that dictates cargo sorting or directional EV release have not been identified. The present study demonstrates that the RPE uses a combination of ESCRT-dependent (Hrs) and ESCRT-independent (ceramide and syntenin-1) mechanisms for EV biogenesis and cargo sorting. These data help establish a stepwise mechanism whereby ceramide, Hrs, syntenin-1, and Rab37 promote the biogenesis of small uniform ILVs containing Cx43. These ILVs are released as EVs likely by the concerted actions of Rabs 11a, 27a, and 37, which are known to function in apical exocytosis and/or EV secretion (18, 65).

As the most abundant gap junction protein in the RPE, Cx43 regulates intercellular communication within the RPE monolayer and with the neuroretina (66, 67). Studies in other tissues have identified non-gap junction roles for Cx43 in ciliogenesis, cytoskeleton remodeling, and cell migration (60, 68–71). Aberrant sorting of Cx43 into ILVs occurs in response to stress or disease (47), and these Cx43-EVs have been shown to promote senescence, inflammation, and influence cell fate in diverse disease models (45, 72). Altered Cx43 trafficking in the RPE could have at least two significant consequences for retinal homeostasis: first, increased intracellular Cx43 could remodel the actin cytoskeleton, which would impact RPE cell structure as evidenced by our morphometric data; and second, Cx43-EVs released by diseased RPE could induce microglial migration as evidenced by the enrichment of Cx43 in subretinal microglia in *Abca4^-/-^* mice. Supporting this hypothesis, iPSC-RPE established from AMD donors have been reported to release EVs containing ceramide and Cx43, and exposure of healthy iPSC-RPE cultures or retinal organoids to these EVs induced cytoskeletal rearrangements, cell fate changes, and immune cell activation in recipient cells (28).

Intracellular ceramide levels are tightly regulated by the concerted actions of ceramide synthases and sphingomyelinases, which generate ceramide, and acid ceramidase, which degrades it (73). Sphingomyelinases are characterized by the pH optima of their hydrolytic activities as acid (ASM) or neutral sphingomyelinases (nSMase). Inhibition of nSMase2 has most often been used to modulate ceramide levels and hence EV biogenesis in various cell types (20, 74–76). However, this is likely to be a cell type-specific effect as nSMase2 inhibition does not alter EV biogenesis in macrophages, cortical neurons, melanoma cells, among others (77, 78). Moreover, inhibiting nSMase2 is associated with altering post-Golgi trafficking and increasing EV release from the cell surface, which could lead to confounding results. We chose to target ASM based on our observation of ASM stabilization and overactivation in Stargardt disease RPE (14–16, 79). Our studies show that inhibiting ASM using the FDA-approved drug desipramine decreases EV formation both *in vitro* and in *Abca4^-/-^* mice RPE. To our knowledge, this is the first evidence that ILV biogenesis in the RPE can be modulated *in vivo* by small molecule drugs delivered systemically. These data also highlight the cell type-specificity of pathological ceramide generation pathways and suggest that ASM overactivation is likely to be the predominant mechanism in the RPE.

In this study, the defined spatiotemporal development of disease pathology in *Abca4^-/-^* mice enabled us to identify ceramide-induced alterations in specific membrane trafficking pathways as an early driver that increases the biogenesis and apical release of Cx43-containing EVs. This, in turn, compromises RPE structural integrity and promotes immune cell activation and photoreceptor functional deficits. The significant challenges associated with tracking the formation and function of RPE-derived EVs *in vivo* have hampered our insight into these enigmatic vesicles in retinal health and disease. The data presented in this study provide a framework to further interrogate the cellular machinery that participates in EV biogenesis and secretion, which will help establish how these contribute to pathology.

Our study reinforces the RPE as a central communication hub in the retina and suggests that therapeutic approaches such as ASM inhibition, which restores membrane trafficking in diseased RPE, could help maintain intra- and intercellular communication and prevent photoreceptor functional deficits in Stargardt disease. Increased ceramide and endosomal deficits in the RPE are common features of AMD and clinically heterogeneous IRDs such as Bietti’s crystalline dystrophy, Batten disease, retinitis pigmentosa, Farber disease, and autosomal dominant Stargardt disease (STGD3) (15, 28, 80–85), indicating a broader relevance for the pathogenic mechanisms and therapeutic targets identified here.

## Acknowledgements

We thank members of the Lakkaraju lab for helpful discussions and D. Michiel Pegtel for the CD63-pHluorin plasmid. This work was supported by NIH grants NEI R01EY023299 (A.L.), NEI R01EY030668 (A.L.), NEI R01EY035514 (A.L.), NEI P30EY002162/EY037668 (Core Grant for Vision Research, UCSF), NEI P30EY016665 (Core Grant for Vision Research, University of Wisconsin, Madison), the BrightFocus Foundation Lorraine Maresca award for Innovative Research in AMD M2021020I (A.L.), Reeves Foundation award for AMD (A.L.), All May See Foundation Postdoctoral Grant Award (L.X.T), National Eye Institute Supplement R01EY030668S1 (N.L.C.) and an unrestricted grant from the Research to Prevent Blindness Foundation (UCSF Department of Ophthalmology).

## Experimental Methods

### Mice

*Abca4^−/−^* (Jackson Labs Abca4tm1Ght/J) and corresponding wildtype mice (Jackson Labs; 129S1/SvlmJ), both on Rpe65 Leu450 background, were raised under 12-h cyclic light with standard diet. For pharmacological studies, mice were administered desipramine (10 mg/kg three times per week for 8 weeks) by intraperitoneal injections Heterozygous P23H knock-in mice (Rho*^P23H/+^*) and corresponding C57Bl/6J wildtype mice were obtained from Jackson labs. Randomized numbers of male and female mice were used in all studies. All studies were approved by the University of California, San Francisco institutional animal care and use committee.

### Polarized primary RPE cultures

RPE were isolated from porcine retinas using established protocols (23). Briefly, the anterior segment was removed at the *ora serrata*, and the retina was gently detached by clipping at the optic nerve head. RPE cells were isolated from eyecups upon incubation with 0.5% trypsin (Lonza, Walkersville, MD) with 5.3 mM EDTA in Hank’s balanced salt solution (Corning, Corning, NY) and plated onto T25 flasks in growth medium (DMEM with 4.5 g/L glucose, 1% heat-inactivated fetal bovine serum, 1% non-essential amino acids and 1% penicillin/streptomycin). After two weeks, cells were trypsinized and plated at confluence (∼ 300,000 cells/cm^2^) on collagen-coated Transwell filters as described (14, 23). Monolayers were treated with the lipofuscin bisretinoid A2E (10 µM for 6 h, 48 h chase) or with U18666A (1 µM, 16 h, Cayman) as previously published (16, 79).

### Immunoblotting

RPE collected from mouse eyecups, polarized primary RPE cultures, or purified EVs were lysed in 1x HNTG buffer supplemented with a protease inhibitor cocktail and resolved by immunoblotting as previously described (86). Samples were resolved on a NuPAGE 4-12% Bis-Tris gel (Invitrogen) and transferred onto nitrocellulose membranes. After blocking, membranes were incubated with indicated primary antibodies diluted in blocking buffer with 0.1% Tween20. Membranes were then incubated with IRDye secondary antibodies (1:5000, LI-COR) for 1 h at room temperature and scanned using LI-COR Odyssey CLx. Band intensities were quantified using Image Studio (LI-COR Odyssey).

### Immunofluorescence staining

Mouse RPE flatmounts or polarized RPE monolayers grown on Transwell filters were fixed in PFA, blocked, and stained with specific primary antibodies (see Table S1 for antibody sources and dilutions) as previously described (86). After incubation with AlexaFluor secondary antibodies (1:500, ThermoFisher Scientific, Waltham, MA), RPE were labeled with phalloidin (1:200, Cytoskeleton, PDHG1) and DAPI (Sigma-Aldrich, St. Louis, MO; D9542, 1:200 in PBS). Mouse flatmounts were treated with TrueBlack (Biotium) for 30 seconds to quench RPE lipofuscin autofluorescence. After rinsing, flatmounts were mounted and sealed on clean slides with PBS and Vectashield Vibrance (Vector Labs, Peterborough, UK).

### Imaging and image analysis

RPE monolayers and mouse flatmounts were imaged using the Nikon spinning disk confocal microscope with 100x (1.49 NA) objective equipped with a Live-SR super-resolution component. The same laser power and exposure time was used for each antibody within a set of experiments. Acquired images were subjected to background subtraction and Gaussian smoothing in Imaris (Bitplane, South Windsor, CT). Where indicated, the image is subjected to automatic 3D deconvolution using NIS-Elements (Nikon, Tokyo, Japan). 3D surface and volumetric reconstructions of Cx43, Iba1/microglia, and CD63 were done using the “Surfaces” tool with the same threshold applied. to all images within an experiment (86, 87). The “Spots” tool was used for quantifying microglia and manual correction was performed as necessary.

### Extracellular vesicle (EV) isolation and purification

Polarized RPE monolayers grown on Transwell filters were cultured in EV-depleted media (27) for 48 h prior to EV harvest. EVs from apical and basal media were isolated via differential ultracentrifugation or size exclusion chromatography according to established protocols (26, 27). For differential ultracentrifugation, media was centrifuged at 300x*g* for 10 minutes to remove cellular contaminants at 4°C in an Eppendorf 5240 centrifuge. Pellets were discarded and supernatants centrifuged in fresh tubes at 2,000x*g* for 10 minutes at 4°C in an Eppendorf 5240 centrifuge. Supernatants were transferred into polyallomer tubes and centrifuged at 10,000xg for 30 minutes at 4°C in a Beckman L8-80M preparative ultracentrifuge with a SW28 rotor to remove apoptotic particles. Supernatants were centrifuged at 100,000x*g* for 3 hours at 4°C to isolate purified EVs. Pellets were rinsed in sterile PBS and centrifuged in an SW44Ti rotor for 100 minutes at 4°C at 100,00 x *g* to remove any external proteins and debris. For size exclusion chromatography, Izon columns (qEV2 35nm, Izon) were used according to manufacturer’s instructions. Briefly, media was concentrated using pre-rinsed Centricon 70-100 kDa filters and centrifuged to remove aggregates at 10,000x*g* for 10 minutes at 4°C. Pellets, if any, were discarded. Supernatants were applied to Izon columns and fractions 6-9 were collected as pure EVs.

### Transmission Electron Microscopy of EVs

TEM of EVs was performed according to published protocols (26, 27). Briefly, purified exosomes were fixed in 4% PFA for 20 minutes before being adsorbed onto a 200-mesh copper slot grid with a carbon-formvar coating. Grids were fixed with 2% glutaraldehyde, rinsed in deionized water, and stained with 4% aqueous uranyl acetate and uranyl oxalate. Grids were imaged using a Philips CM120 TEM at an accelerating voltage of 80 kV and 2 µm spot size using a BioSprint 12 camera (AMT) at a 1 s exposure and 2-frame average. The entire grid was imaged. Images were analyzed using the AMT plugin for FIJI to calibrate pixel sizes.

### NanoTracking Analysis of EVs

Purified EVs were resuspended in PBS that was pre-filtered through a 20 nm pore size fiberglass filter (Anotop 6809-1002, GE) and measured using an LM10 NanoSight (Malvern). Five 30-second videos were acquired per sample, and each sample was measured in triplicate. Camera setting of 13 with a sensitivity threshold of 3 was used for both acquisition and analysis. Between each measurement, the optical flat and instrument was thoroughly cleaned with pure ethanol and deionized water. A background video was taken of the filtered water to subtract contaminating particles.

### Live imaging of EV release

Primary RPE were transfected with CD63-pHluorin (5 µg DNA per 1.5×10^6^ cells, Lonza Nucleofector) (35) and plated onto serum-coated glass-bottom dishes. Cell monolayers were treated with A2E alone (10 µM for 6 h followed by a 48 h chase in fresh culture medium) or with A2E and desipramine (10 µM, 3 h) (14–16, 23). High-resolution high-speed live imaging was performed at 37°C in an Okolab humidified microenvironmental chamber on the Nikon spinning disk confocal microscope using a CFI60 Apochromat TIRF 100X oil immersion objective (1.49 NA) as described previously (79, 86, 87). Identical laser power, exposure and gain settings were applied for all conditions within a set of experiments. To capture apical exocytosis, planes with pre-docked CD63-pHluorin vesicles at the apical membrane were identified and time-lapse movies were acquired at 30 s intervals for 10 min (50 msec exposure). Surface reconstructions were performed with the “Surfaces” module with tracking enabled on Imaris. The appearance of new surfaces absent in the preceding time frame were quantified as exocytic events.

### Super-resolution imaging of single EVs

Distribution of Cx43 and ceramide in single EVs was analyzed by super-resolution imaging using dSTORM system equipped with a 100X 1.45 NA oil immersion objective (ONI Nanoimager, Oxford Nanoimaging, UK). EVs purified by size exclusion chromatography were stained overnight with FITC-conjugated CD9 antibody (EV Profiler kit, ONI) and antibodies to Cx43 and Ceramide (conjugated to 561 and 647 dyes, respectively, Biotium Mix-n-Stain kits). Samples were immobilized on the ONI chip and imaged using ONI’s B-Cubed buffer. Approximately 1,500 frames were acquired at a 30° angle with a laser power of 15%. The standard imaging protocol was applied along with CODI drift correction. Colocalization analysis of specific molecules in a single EV was performed using the CODI online analysis platform (https://alto.codi.bio/). Images were further analyzed using Imaris with intensity-based colocalization and background subtraction and Gaussian filtering.

### Proteomic analyses of EVs

RPE cultures treated with vehicle or U18666A (2.5 µM, 16 hours) were cultured in exosome-depleted medium overnight. EVs were purified from the media by differential ultracentrifugation as described above. The resultant pellet was resuspended in PBS and purified using a continuous iodixanol gradient. The gradient was then harvested in 200 µl increments and absorption spectra were matched to determine the ideal percentage iodixanol for EV purification. This EV fraction was subjected to LC-MS/MS using an Orbitrap Elite mass spectrometer (ThermoFisher Scientific) connected to an 1100 Nano flow HPLC system (Agilent) with temperature-controlled autosampler and temperature-controlled column compartment (Mass Spectrometry Core at the University of Wisconsin-Madison). Samples were separated by HPLC using EasySpray 75 µm columns coupled to an EasySpray source (ThermoFisher). Data were analyzed using Proteome Discoverer (ThermoFisher), MASCOT, and Scaffold. Samples were analyzed in duplicate. A false discovery rate (FDR) of 1% was used as threshold for both protein and peptide levels. Protein-protein interaction (PPI) network and gene ontology (GO) enrichment analyses were performed using STRING.

### Analyses of EVs from MDCK cultures

Type II MDCK cells (ATCC) were cultured on Transwell filters as described (88). EV isolation, purification, and analyses were performed as above.

### Transmission Electron Microscopy of mouse retinas

Enucleated whole globes were fixed in 4% PFA, 5% glutaraldehyde in 0.1M Sodium Cacodylate pH 7.4 for 1 hour at room temperature. The anterior portion was then removed (cornea, lens, iris) and relaxing cuts were made to flatten the eyes, with the retina remaining attached. Eyes were then rinsed in 0.1 M Sodium Cacodylate pH 7.4 three times before being moved into 1-Hexadecene. Five 2 mm punches were taken from each eye with a biopsy punch (Integra). These punches were then moved into 0.3 mm Type B freezing hats (Ted Pella) coated in 1-Hexadecene. Excess 1-Hexadecene was applied to remove air before being capped with another Type B freezing hat for a total space of 0.6 mm. These were subjected to high-pressure freezing in a Balzers HPM-010. Hats were separated under liquid nitrogen and placed in frozen freeze substitution media (0.1% methanolic uranyl acetate and 4% osmium tetroxide in pure acetone). The freeze substitution media was placed into a Leica AFS set to - 70°C for 72 h before ramping up to 4°C over the course of 24 h. Excess freeze substitution media was removed by three rinses in pure acetone. Hats were removed and samples were incubated with mixtures of 50:50 and 75:25 complete Embed-812 Hard formulation/acetone for 16 hours each with rotation. Following this, three changes of pure resin were conducted over the course of eight hours before being held under vacuum overnight. Samples were removed and put into fresh resin in coffin molds with labels. Sections were cut using an ultramicrotome (Leica) at 70 nm thickness and placed onto two 1 mm copper slot grids with formvar. Grids were stained using 4% uranyl acetate in a Hiroka staining chamber, rinsed with deionized water and stained with lead citrate in a CO_2_ depleted environment created by sodium hydroxide pellets. Grids were dried and imaged on a Philips CM120 at an accelerating voltage of 80 kV and 2 µm spot size using a BioSprint 12 camera (AMT) at a 1 s exposure and 2-frame average. The entire grid was imaged.

Images were analyzed using the AMT plugin for FIJI to calibrate pixel sizes. ILVs were identified based on the presence of a complete membrane surrounding the vesicle and a clear doughnut morphology. For an organelle to be counted as a MVB required at least 2 ILVs. Diameters of ILVs and MVBs were measured using the Line tool. To classify MVBs as apical or basolateral, low magnification shots of the RPE were taken to encompass basal infoldings and microvilli. Distances were measured from the bottom of the basal infoldings to the base of the microvilli. MVB polarity was determined based on whether the MVB was within 2 µm of the apical or basolateral membrane of the RPE. To calculate MVBs per micron, a “burn” was performed on the grid by increasing the beam intensity to make a mark at the start of each imaging session and at the end. The lowest mag possible on the scope was then used to image the entire grid. The Segment tool in FIJI with the AMT plugin was used to trace along the basolateral infoldings of the RPE from each burn. This total distance was used to calculate MVBs per micron.

### Segmentation and morphometric analyses of the RPE monolayer

Individual channels from de-identified 10x stitched images of mouse RPE flatmounts were split and max-intensity projections of the phalloidin channel were generated using a custom-built macro code in Fiji. Then, whole flatmounts were segmented using the machine-learning-based algorithm Cellpose 4.0.5 (89) with Python 3.12.7. The base model and the following parameters were used to successfully segment RPE cells across conditions: diameter=blank, flow threshold=0.5, cellprob threshold=-6.0, lower=1.0, upper=99 and niter dynamics=1600. Using the GUI, only correctly segmented cells were saved and exported as regions of interest (ROI) for further shape analysis in Fiji. Using a custom-built code in Fiji, max-intensity projected images were opened with their respective ROI to obtain cell area and aspect ratio measurements using the built-in *measure* tool in Fiji. The results were stored in an Excel sheet and re-identified to their respective groups and conditions. To generate color-coded maps, a Python 3.12.7 script was used to convert Cellpose-generated masks into arrays, enabling color segmentations based on cell area. The resulting measurements were processed in Prism 10 using one-way ANOVA tests with Tukey’s multiple comparisons.

### Electroretinography

ERGs were recorded using the Celeris rodent electrophysiology system (Diagnosys LLC, Lowell, MA). Mice were dark adapted overnight and anesthetized intraperitoneally with ketamine (80 mg/kg) and xylazine (10 mg/kg) under dim red illumination. Whiskers were trimmed and a mydriatic solution composed of 5% phenylephrine and 1% tropicamide was applied to the eyes. Generous amounts of 0.3% hypromellose gel (Genteal) were applied to keep the eyes moist. The Celeris system was used with bright stimulators in an active/reference/ground configuration. The grounding electrode needle was placed into the skin of the animal near the tail. The reference electrode needle was placed in the cheek of the animal and the warming pad was set to 37℃. An 8-step series was taken with 10 sweeps per step with each sweep lasting four seconds with 20 seconds between each sweep (90). Sweeps were averaged together for a final readout with outlier readouts removed. Brightness was set from 0.01 to 25 cd·s/m². After the recording, atipamezole was administered and mice were allowed to recover in a warm clean cage. Wave peaks and times were determined by Espion V6 (Diagnosys) software with a-waves starting at 10 ms after stimulus with a 30 ms range, b-waves had a start time 25 ms after stimulus with a 60 ms range, and c-waves had a peak start time at 300 ms with a 3,000 ms range. The peak amplitudes and times from each eye were averaged together and analyzed in Prism using two-way ANOVA with Bonferroni’s multiple comparisons test.

### Statistics

Data were analyzed by unpaired two-tailed *t*-test with Welch’s correction for two experimental groups, and one-or two-way ANOVA tests with post-hoc analysis for comparison between multiple groups (Graphpad Prism). Data are presented as Mean ± SEM, with ≥ 3 mice per age, genotype, or treatment.

## Author contributions

CJG, SW, LXT, and AL designed experiments, analyzed data, and wrote the manuscript.

CJG, SW, VF, REL, NLC, LXT, and SRP performed experiments and analyzed data.

EO and FME provided new reagents and access to instrumentation.

AL designed the study and provided funding.

All authors edited the manuscript.

