## Supplementary figures, legends, methods for "Acid sphingomyelinase inhibition restores RPE homeostasis and photoreceptor function in preclinical Stargardt macular degeneration models"

**This PDF file includes:**

Figures S1 to S4 with legends

Table S1. Proteomic analyses of EVs

Table S2. Key resources

Detailed Experimental Methods

Supplementary References

Figure S1

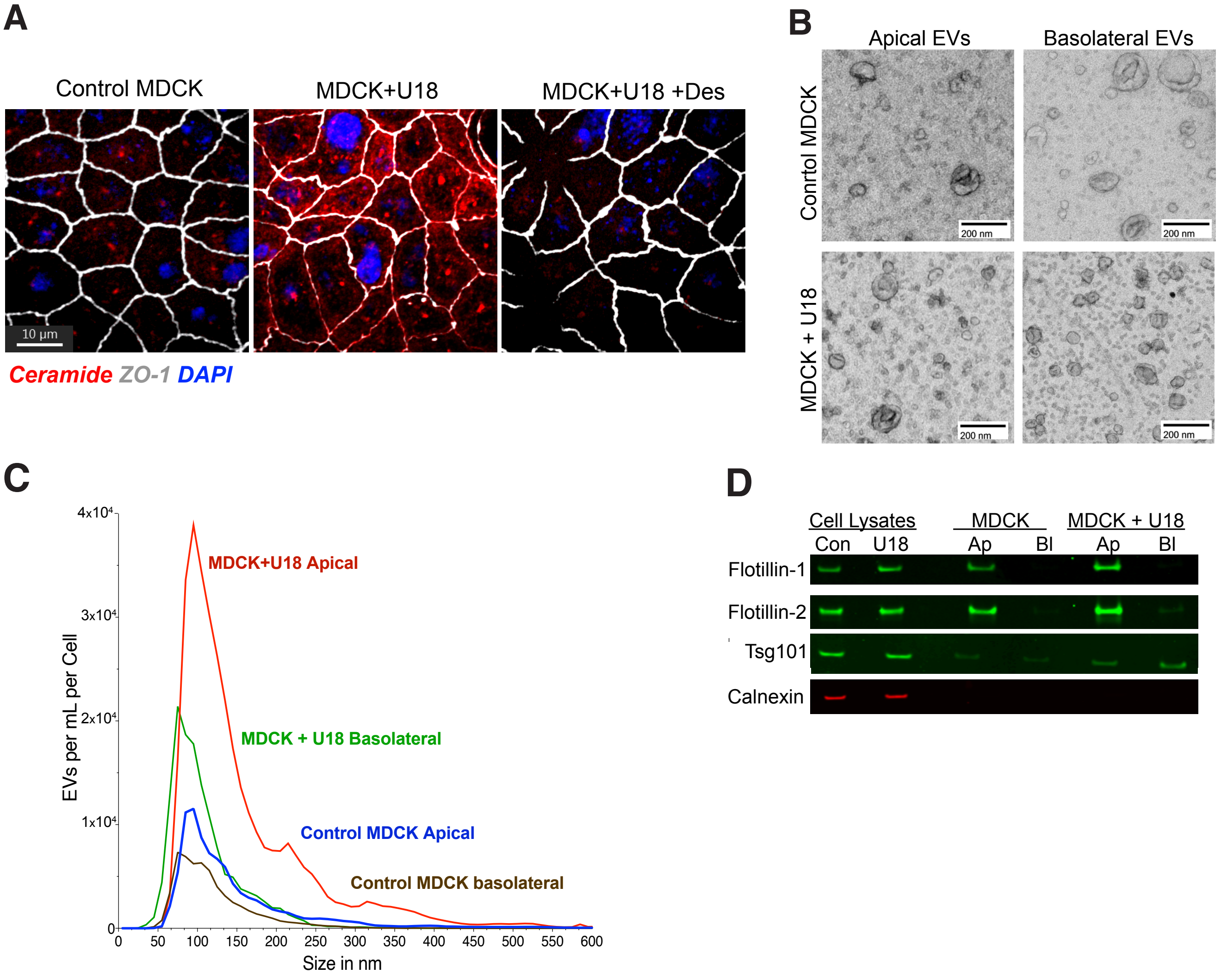

**Figure S1. Ceramide drives apical EV secretion from polarized MDCK cultures. (A)** Immunostaining of ceramide (red), ZO-1 (white) and DAPI (blue) in polarized MDCK monolayers untreated, or treated with U18666A (U18) and desipramine (des). **(B)** Representative electron micrographs of EVs released into the apical or basal media of MDCK cultures ± U18. **(C)** Number and size distribution of EVs released by polarized MDCK ± U18. **(D)** Representative immunoblot of total cell lysates and apical (Ap) or basolaterally (Bl) released EVs from MDCK cultures ± U18.

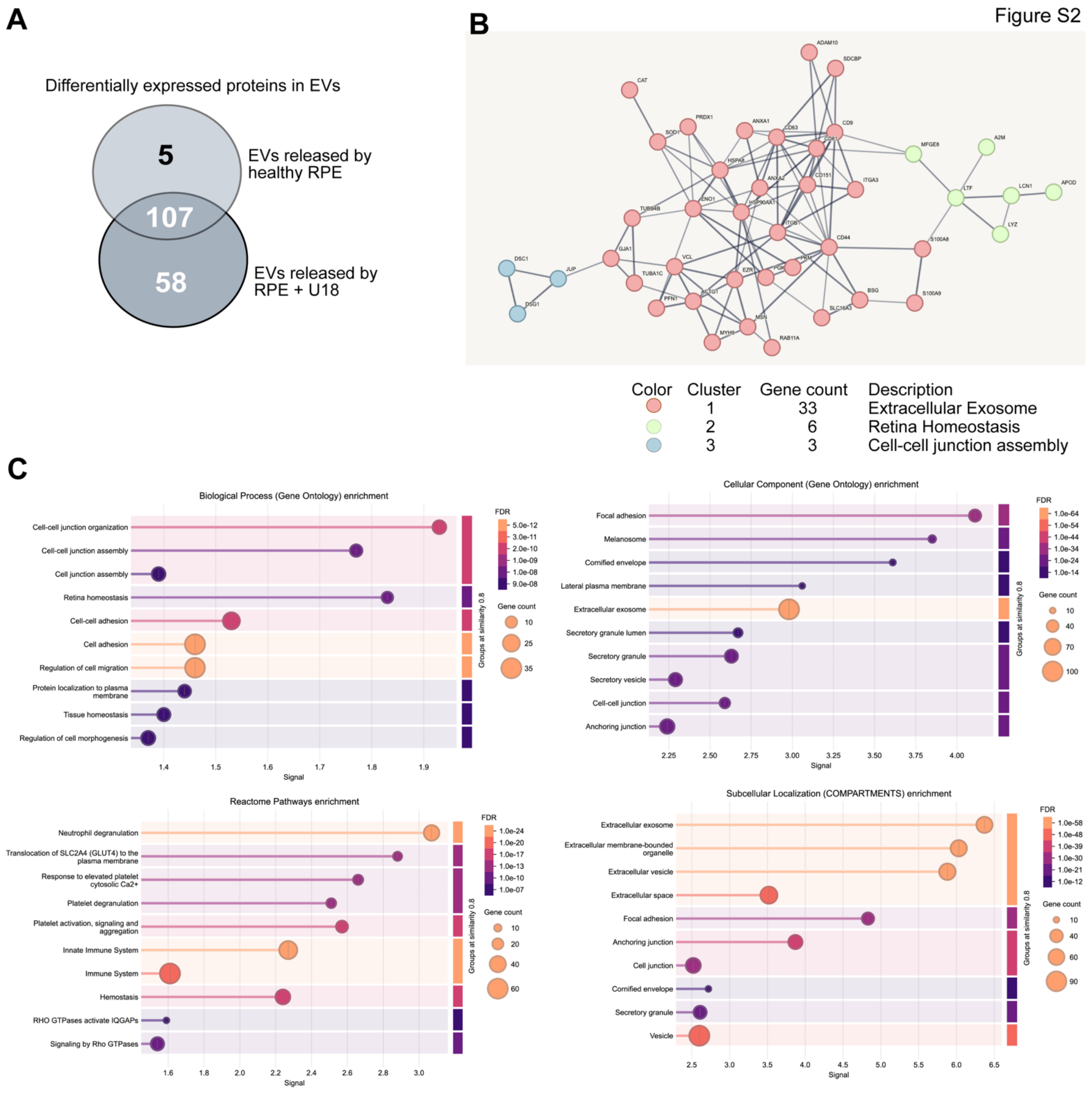

**Figure S2. LC-MS/MS analysis of EV proteins. (A)** Venn diagram showing the number of differentially expressed proteins in each population of EVs. **(B)** STRING network analysis of protein-protein interactions (PPI) of proteins identified in EVs and top 3 clusters of protein nodes calculated by k-means. **(C)** Gene ontology (GO) and KEGG reactome analyses of EV proteins based on biological process, cellular component, and subcellular localization.

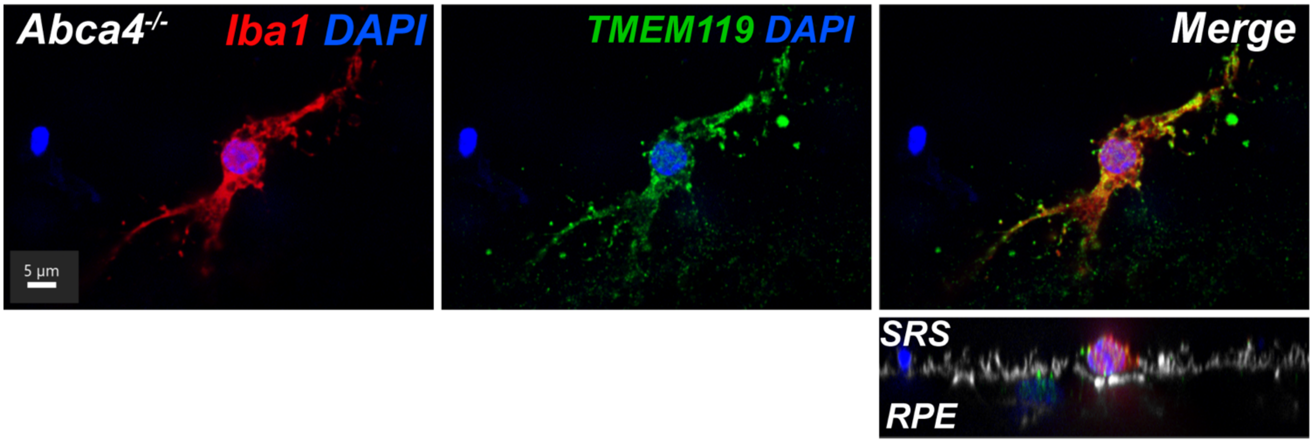

**Figure S3.** Immune cells in the subretinal space of 18-mo *Abca4*$^{-/-}$ mice RPE express canonical microglial markers. Microglia stained for Iba1 (red) and TMEM119 (green). The x-z image shows microglia in the subretinal space (SRS) sitting atop RPE microvilli stained with phalloidin (gray).

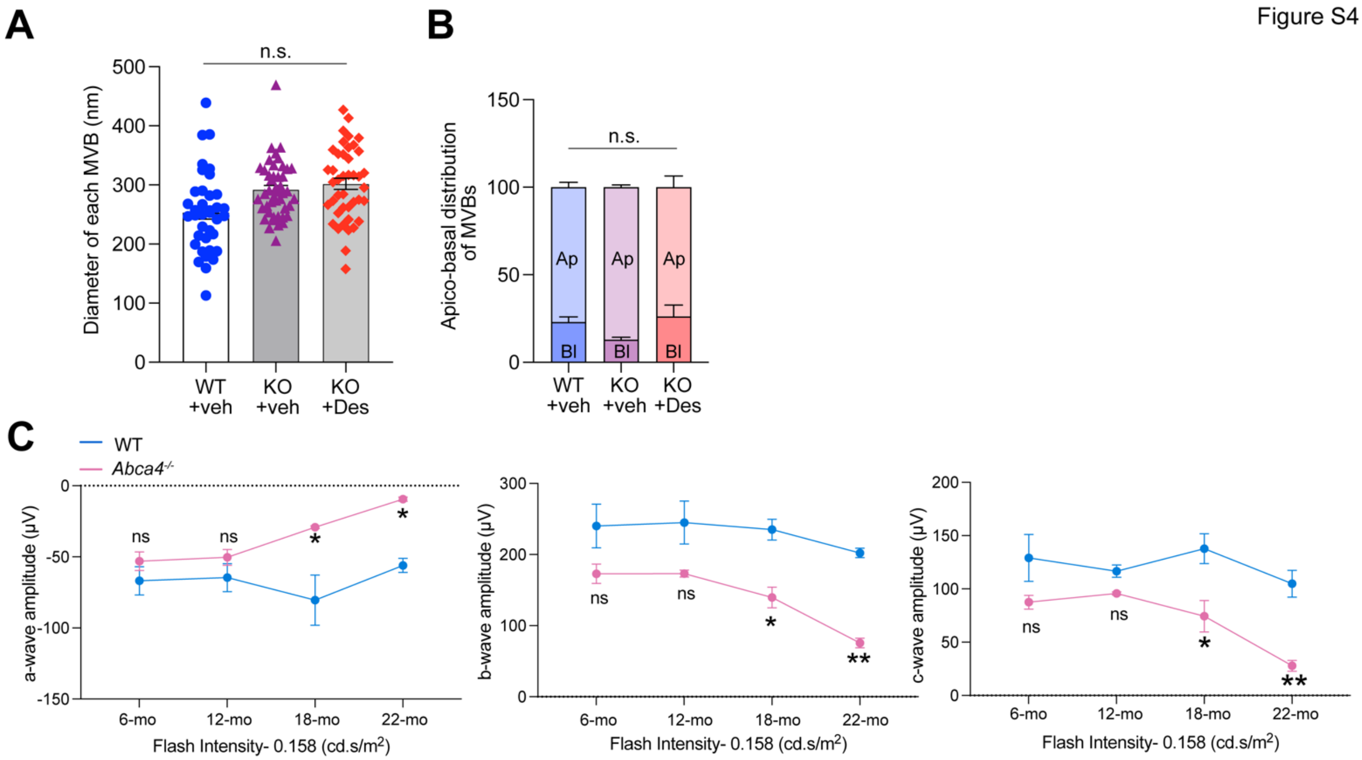

**Figure S4. (A)** Diameter and **(B)** apico-basal distribution of MVBs in 18-month-old wildtype and *Abca4*$^{-/-}$ mice treated with vehicle or desipramine analyzed from transmission electron micrographs. n.s. not significant. **(C)** Scotopic a-, b-, and c-waves in 6, 12, 18, and 22-month-old wildtype and *Abca4*$^{-/-}$ mice measured at 0.158 cd.s/m$^2$. Mean ± SEM, n = 3-4 mice/genotype and age. *, $p < 0.05$ and **, $p < 0.01$ by two-way ANOVA and Tukey's multiple comparisons test.

**Table S1. Proteins enriched in EVs released by RPE treated with U18**

| Protein | Gene Name | Fold Enrichment in EVs from RPE+U18 |
|---|---|---|
| Connexin 43 | GJA1 | 1158.70 |
| CD147 | BSG | 850.39 |
| Phosphoglycerate kinase 1 | PGK1 | 792.56 |
| Arf3 | ARF3 | 704.05 |
| Annexin A3 | ANXA3 | 666.74 |
| CD63 | CD63 | 657.14 |
| Moesin | MSN | 648.76 |
| Rab10 | RAB10 | 630.85 |
| Rab13 | RAB13 | 623.60 |
| Superoxide dismutase 1 | SOD1 | 578.66 |
| Syntenin-1 | SDCBP | 534.90 |
| Heat shock protein 70 | HSP70 | 462.43 |
| Beta-tubulin | TUBB4B | 461.70 |
| Rab11 | RAB11A | 393.00 |
| Ezrin | EZR | 391.20 |
| MCT4 | SLC16A3 | 360.81 |
| CD151 | CD151 | 307.54 |
| Hrs | HGS | 293.73 |
| Alix | PDCD6IP | 272.69 |
| Hsp90 | HSP90AA1 | 256.17 |
| Lactadherin | MFGE8 | 194.94 |
| Profilin | PFN1 | 153.11 |
| Alpha-tubulin | TUBA1C | 143.50 |
| Adam10 | ADAM10 | 97.22 |
| Integrin b1 | ITGB1 | 13.80 |
| Calmodulin | CALM2 | 12.92 |
| CD44 | CD44 | 12.69 |
| Hsc71 | HSPA8 | 11.27 |
| Integrin alpha3 | ITGA3 | 9.81 |
| Vinculin | VCL | 9.35 |
| Annexin A1 | ANXA1 | 7.58 |
| Pyruvate kinase | PKM | 7.05 |
| Annexin A2 | ANXA2 | 5.55 |
| Myosin 9 | MYH9 | 4.44 |
| Catalase | CAT | 4.00 |
| S100-A9 | S100A9 | 3.77 |
| CD9 | CD9 | 3.20 |
| Actin | ACTG1 | 3.09 |
| Alpha-enolase | ENO1 | 2.60 |
| Peroxiredoxin-1 | PRDX1 | 2.56 |
| S100-A8 | S100A8 | 2.28 |
| Junction plakoglobin | JUP | 2.00 |
| Desmocollin-1 | DSC1 | 1.50 |
| CD81 | CD81 | 1.46 |
| Fatty acid binding protein 5 | FABP5 | 1.46 |
| Lipocalin-1 | LCN1 | 1.46 |
| Apolipoprotein D | APOD | 1.46 |
| Desmoglein-1 | DSG1 | 1.40 |
| Lysozyme | LYZ | 0.37 |
| Lactotransferrin | LTF | 0.34 |
| Alpha-2 macroglobulin | A2M | 0.03 |

**Table S2. Key Resources Table**

| **Antibody** | **Source** | **Catalog Number** | **IF** | **WB** | **Host** |
|---|---|---|---|---|---|
| Acid Sphingomyelinase | Abcam | Ab83554 | 1:100 | | Rabbit |
| ActiStain | Cytoskeleton | PHDG1-A | 1:200 | | |
| Alix | Abcam | Ab50249 | | 1:500 | Rabbit |
| Calnexin | Abcam | Ab75801 | | 1:2000 | Rabbit |
| CD63 | LSBio | LS-C204227 | 1:100 | | Goat |
| Ceramide | Enzo | 86153-838 | 1:10 | | Mouse |
| Connexin 43 | Abcam | Ab11370 | 1:200 | | Rabbit |
| Connexin 43 | Abcam | Ab79010 | 1:100 | | Mouse |
| Flotillin-2 | Millipore Sigma | SAB2500405 | | 1:500 | Goat |
| Hrs | Novus | NB1002839 | 1:200 | 1:500 | Goat |
| Hsp70 | BD Biosciences | 610608 | | 1:1000 | Mouse |
| Iba1 | Wako | 019-19741 | 1:200 | | Rabbit |
| Rab7 | Cell Signaling | 9367S | | 1:1000 | Rabbit |
| Rab8 | BD | 610845 | | 1:1000 | Mouse |
| Rab10 | Abcam | 1014853 | | 1:1000 | Rabbit |
| Rab11a | Cell Signaling | 5589S | 1:50 | 1:1000 | Rabbit |
| Rab27a | Proteintech | 17817-1-AP | 1:100 | 1:1000 | Rabbit |
| Rab37 | Novus | NBP2-20042 | 1:100 | 1:1000 | Rabbit |
| Rhodopsin | Millipore | MABN15 | 1:200 | | Mouse |
| RPE65 | Novus | NB100-355 | | 1:1000 | Mouse |
| IRDye | LI-COR | | | 1:5000 | |
| DC Assay Kit | BioRad | NC9372425 | | | |
| NuPAGE 1.5mm 4-12% Bis-Tris Gel | Invitrogen | NP0336BOX | | | |
| **Assay Reagents** | **Source** | **Catalog Number** | | | |
| Protease Inhibitor Cocktail | EMD Millipore | 539134-1ML | | | |
| Phosphatase Inhibitor Cocktail | Fisher | PI78420 | | | |
| NuPAGE 4xLDS Sample Buffer | Invitrogen | NP0007 | | | |
| SeeBlue Plus 2 Ladder | Invitrogen | LC5925 | | | |
| NuPAGE 10x Reducing Agent | Invitrogen | NP0009 | | | |
| Antioxidant | Invitrogen | NP0005 | | | |
| Embed-812 Kit | Electron Microscopy Sciences | 14121 | | | |
| Integra Biopsy Punches | Milltex | 33-31 | | | |
| Type B Freezing Hats | Pelco | 50-193-1254 | | | |

| | | | |
|---|---|---|---|
| 2mm Copper Slot Grids | Pelco | 4514 | |
| 200 Mesh Copper Grids | Pelco | 01800-F | |
| TissueTek OCT | Sakura Finetek | 25608-930 | |
| EV Profiler Kit | ONI | 900-00208 | |
| Mix-n-Stain Antibody Labeling Kit 561 | Biotium | 89171-586 (EA) | |
| Mix-n-Stain 647 CF | Biotium | 89171-594 (EA) | |
| AlexaFluor Secondary (various colors) | ThermoFisher | | 1:500 |
| IRDye Secondary | LI-COR | | 1:10,000 |
| **Plasmid** | **Source** | **Reference** | |
| CD63-pHlourin Plasmid | Gift from Dr. Pegtel's lab | Verweij et al., 2018 | |
| **Drugs** | **Source** | **Catalog Number** | |
| Desipramine | Enzo | BML-AR119-0100 | |
| Phenylepherine | Acros Organics | 20724010 | |
| Tropicamide | Alfa Aesar | J61132 | |
| **Cell Sources** | | | |
| Primary porcine RPE | Established from freshly harvested pig eyes obtained from Hart & Vold Meat Market, Baraboo, WI (*1*) and Lampire Biologicals, Pipersville, PA. | | |
| MDCK Type II | ATCC | CRL-2936 | |
| **Chemicals** | **Source** | **Catalog Number** | |
| Bovine Serum Albumin | Rockland Immunochemicals | BSA-50 | |
| Calcium Chloride Dihydrate | Sigma-Aldrich | C7902 | |
| DAPI (14.3 mM stock solution) | Sigma-Aldrich | D9542 | 1:200 |
| Glucose | Sigma Aldrich | G7528 | |
| HBSS | Corning | 21-023-CV | |
| HEPES | ThermoFisher | 15630080 | |

| | | | |
|---|---|---|---|
| Magnesium Chloride Hexahydrate | Sigma-Aldrich | M2393 | |
| Paraformaldehyde (8%) | Electron Microscopy Sciences | 157-8 | |
| Phosphate Buffer Saline | ThermoFisher | BP665-1 | |
| Glutaraldehyde (25%) | Electron Microscopy Sciences | 16220 | |
| 1-Hexadecene | TLC | H0323 | |
| Methylcellulose | Sigma-Aldrich | M6385-100G | |
| Uranyl Acetate | Electron Microscopy Sciences | 22400 | |
| Lead Citrate Trihydrate Air-Free | Electron Microscopy Sciences | 22410 | |
| Sodium Cacodylate Trihydrate | Electron Microscopy Sciences | 12300 | |
| Osmium Tetroxide | Electron Microscopy Sciences | | |
| Pure Acetone | Electron Microscopy Sciences | 10000 | |
| 200 Proof Ethanol | Koptec | V1001 | |
| Saponin | Sigma-Aldrich | 84510 | |
| Sucrose | Sigma-Aldrich | S0389 | |
| TrueBlack | Biotium | 23007 | 1:20 |
| VectaShield | Vector Laboratories | H1000 | |
| Ciprofloxacin | Sigma-Aldrich | 17850-5G-F | |
| DMEM | Corning | 10-013-CV | |
| Fetal Bovine Serum (heat-inactivated) | American Type Culture Collection | 30-2020 | |
| Non-essential amino acids (NEAA) | Corning | 25-025-CI | |
| Penicillin-Streptomycin | Corning | 30-002-CI | |
| 0.25% Trypsin | Corning, Corning, NY | 25-053-CI | |
| 2.5% Trypsin | Lonza | 17-160E | |

| | | |
|---|---|---|
| Intercept (PBS) Protein-Free Blocking Buffer | LI-COR | 927-90001 |
| **Software** | | |
| Prism | Graphpad | |
| NIS Elements | Nikon | |
| Imaris | Bitplane | |
| CODI | ONI | https://alto.codi.bio |
| Creative Suite | Adobe | |

### Detailed Experimental Methods

#### Mice
